# PP2A^Cdc55^ dephosphorylates Pds1 to inhibit spindle elongation

**DOI:** 10.1101/648386

**Authors:** Shoily Khondker, Sam Kajjo, Devon Chandler-Brown, Jan Skotheim, Adam Rudner, Amy Ikui

## Abstract

DNA replication stress stalls replication forks leading to chromosome breakage and Intra-S checkpoint activation. In *S. cerevisiae*, this checkpoint arrests the cell cycle by stabilizing securin (Pds1) and inhibiting the cyclin dependent kinase (CDK) through multiple pathways. Pds1 inhibits separase (Esp1) which cleaves the cohesin subunit Scc1 and also functions in spindle elongation. However, the role of Pds1-Esp1 in spindle elongation during replication stress response is unknown. Here, we show that Pds1 phosphorylation plays a positive role in spindle elongation through the Pds1-Esp1 interaction in unperturbed and replication stress conditions. PP2A^Cdc55^ directly dephosphorylates Pds1 both *in vivo* and *in vitro*. Pds1 hyperphosphorylation in a *cdc55Δ* mutant enhanced the Pds1-Esp1 interaction, which accelerated spindle elongation. This PP2A^Cdc55^-dependent Pds1 dephosphorylation plays a role during replication stress and acts independently of the known Mec1, Swe1 or Spindle Assembly Checkpoint (SAC) checkpoint pathways. We propose a model where PP2A^Cdc55^ dephosphorylates Pds1 to disrupt the Pds1-Esp1 interaction that inhibits spindle elongation during replication stress.

## Introduction

Eukaryotic cells have multiple cell cycle checkpoints to ensure accurate replication and partitioning of genetic information. DNA replication stress compromises genome integrity due to replication fork stalling and collapse which leads to chromosome breakage and DNA damage (*1*). In *S. cerevisiae*, the Intra-S checkpoint is a well-studied pathway that is activated upon replication stress which mainly targets and stabilizes securin (Pds1) (Figure 1A). The sensor kinase Mec1 interacts with Ddc2 to form foci at replication stress sites (*1*, *2*). Mec1 phosphorylates Rad9, which in turn recruits Rad53 to phosphorylate multiple downstream targets. The Rad53 pathway disrupts the interaction between Pds1 and Cdc20, an activator for E3 ubiquitin ligase anaphase promoting complex (APC), leading to Pds1 stabilization and metaphase arrest (*3–8*) (Figure 1A). Furthermore, Rad53 inhibits mitotic cyclin dependent kinase (CDK) activity which also arrests the cell cycle (*9*) (Figure 1A). Mec1 is similarly activated by DNA damage induced by methyl methanesulfonate (MMS), and activates Chk1, another effector kinase, to stabilize Pds1 (*3*, *4*, *10*). Chk1 phosphorylates Pds1 at nine phosphorylation sites to prevent ubiquitination by APC^Cdc20^ (*3–5*). These Chk1 sites are only phosphorylated during MMS-induced DNA damage and are distinct from the CDK consensus sites (*3*). A second replication stress pathway inhibits mitotic CDK through Swe1 phosphorylation on Y19 (*9*, *11*). The Cdc28-Y19 phosphorylation is removed by the phosphatase Mih1 at mitotic entry (*12*). The CDK inhibition by Swe1 is redundant with Rad53*, i.e.,* in the absence of the Intra-S checkpoint, Swe1 prevents sister chromatid segregation during replication stress (*20*). The Spindle Assembly Checkpoint (SAC) is a third pathway important for replication stress and preventing sister chromatid segregation (*9*, *13*) (Figure 1A). The central SAC protein Mad2 physically disrupts binding between the APC and Cdc20 leading to Pds1 stabilization and metaphase arrest (*14–16*).

**Figure 1.**
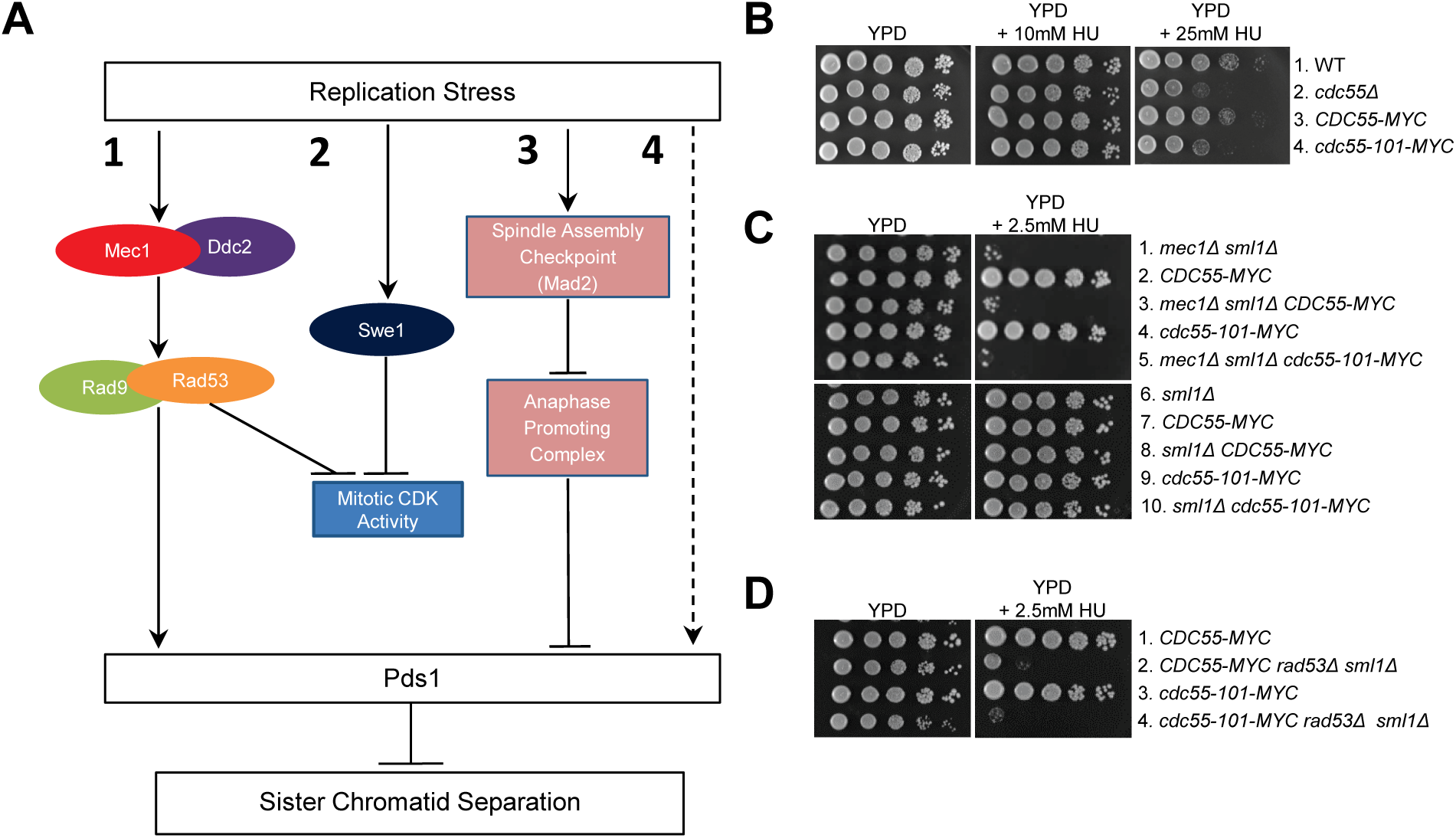
Signal transduction pathways activated by replication stress. (A) 1: The Intra-S checkpoint is triggered by sensor kinase Mec1 and its binding partner Ddc2. Rad53 is an effector kinase downstream of Mec1 which stabilizes Pds1 and inhibits CDK. 2: Swe1 dependent CDK inhibition lead to mitotic arrest. 3: Mad2 is a key protein in SAC which inhibits APC leading to Pds1 stabilization. 4: Unknown pathway. (B-D) Indicated yeast strains were diluted in a 10-fold serial dilution and spotted on YPD plates with HU.

Pds1 is directly regulated by the cyclin-dependent kinase (CDK) Cdc28 through 5 consensus phosphorylation sites (S/TP) during an unperturbed cell cycle. There are two Cdc28 consensus sites located on Pds1 N-terminal region (T27 and S71), which are in close proximity to a KEN Box motif (residues 8-10) and a Destruction-box (D-box) motif (residues 84-93). These sites are necessary for recognition APC^Cdc20^. At the metaphase-to-anaphase transition, Pds1 is targeted for ubiquitination by APC with its activator Cdc20 (APC^Cdc20^) (*15*, *16*). The function and significance of the two N-terminal Cdc28 sites are unknown. The remaining three Cdc28 consensus sites are at the C-terminus (S277, S292, T304) which are necessary for a physical interaction between Pds1 and Esp1 (*17*). The phosphorylated Pds1 recruits Esp1 in the nucleus and on the mitotic spindle (*18*). Esp1 has two independent functions in mitosis: cohesin cleavage as a separase and spindle elongation (*18*). Esp1 is localized to the spindle pole bodies and the spindle midzone during anaphase, and this localization requires Pds1 (*18*). In *pds1Δ* cells, Esp1 does not associate with the spindle (*18*). A temperature sensitive *esp1-C113* mutant, which contains a mutation in the C-terminus in the catalytic domain, fails to elongate spindle in mitosis (*19*). While the role of Esp1 in Scc1 cleavage has been extensively studied, there is less known about its function in spindle elongation.

The serine-threonine phosphatases reverse CDK-dependent phosphorylation. There are two key phosphatases that regulate cell cycle progression: PP2A and Cdc14. PP2A is a heterotrimer consisting of an “A” scaffolding unit (Tpd3), a “B” regulatory unit (either Cdc55 or Rts1), and a “C” catalytic unit (either Pph21 or Pph22) (*20*). PP2A^Cdc55^ and PP2A^Rts1^ have distinct functions and targets (*21*). PP2A^Cdc55^ dephosphorylates CDK substrates and preferentially targets threonine residues (*22*). CDK substrate dephosphorylation occurs late in the cell cycle, and is ordered to ensure the correct sequence of cell cycle events (*22*). PP2A^Cdc55^ is present in both the nucleus and the cytoplasm and its function is dependent on its localization (*23*, *24*). Zds1 and Zds2, two yeast-specific proteins, physically interact with Cdc55 to trap it in the cytoplasm (*24*, *25*). In the cytoplasm, PP2A^Cdc55^ inhibits Swe1 to inhibit mitotic CDK activity and promotes mitotic entry (*11*, *26*, *27*). In the absence of Zds1/Zds2, Cdc55 accumulates in the nucleus (*24*, *25*). In the nucleus, PP2A^Cdc55^ inhibits mitotic progression through two mechanisms: inhibiting APC^Cdc20^ and preventing Cdc14 release (*28*, *29*). Nuclear PP2A^Cdc55^ also prevents the Cdc14 release from nucleolus, a phosphatase that promotes mitotic exit (*29*). Thus, the two phosphatases PP2A^Cdc55^ and Cdc14 play important roles in regulating cell cycle progression through mitosis. PP2A^Cdc55^ plays a role in SAC by dephosphorylating APC components Cdc16 and Cdc27 to inhibit APC activity leading to metaphase arrest (*26*, *30*). While the role of PP2A^Cdc55^ in SAC has been well established, the involvement of PP2A^Cdc55^ in replication stress has been understudied.

The reuse of molecules regulating the metaphase-to-anaphase transition and the cell cycle response to replication stress led us to hypothesize that the same phosphatase may regulate these two processes as well. Consistent with this hypothesis, a previous study showed that *cdc55Δ* was sensitive to hydroxyurea (HU) in a growth assay indicating a possible role of Cdc55 in replication stress (*31*). Here, we investigated if PP2A^Cdc55^ plays a role in replication stress, either as a part of a known checkpoint or through a novel mechanism (*31*). We focused on the nuclear PP2A^Cdc55^ function by using a localization mutant, *cdc55-101*, that excludes Cdc55 from the nucleus (*32*). In this study, we identify a role for PP2A^Cdc55^ during replication stress. PP2A^Cdc55^ dephosphorylates Pds1 both *in vitro* and *in vivo* to inhibit spindle elongation. Our findings show that PP2A^Cdc55^ acts independently of Mec1, Swe1 and SAC to cause robust cell cycle arrest during replication stress.

## Methods

### Plasmids and Strains

Standard methods were used for mating, tetrad dissection, and transformation. HA and MYC tags were generated by standard PCR method (*33*). *PDS1-6HA* was generated using pYM3 plasmid (S3-5’CAGCGAAGAAGGCCTCGATCCTGAAGAACTAGAGGACTTAGTTACTCGTACGCTGCAGGTCGAC 3’; S2-5’CTGTATATACGTGTATATATGTTGTGTGTATGTGAATGAGCAGTGGATATCGATGAATTCGAGCTCG 3’)(*33*). *ESP1-9MYC* was generated using pYM6 plasmid (S3-5’GGCGC AGCTCCTGTTATTTATGGGTTACCGATCAAGTTCGTATCACGTACGCTGCAGGTCGAC 3’; S2-5’CAATGCCTATATGAAATCTTTTCGAAACAGCCAGTACATGTAACAAATCGATGAATTCGAGCTC 3’)(*33*). *CDC16-6HA* was generated using pYM3 plasmid (S3-5’GCCTCGATCCTGAAGAACTAGAGGACTTA GTTACTCGTACGCTGCAGGTCGAC ACGCAGATATGGAACTGGAATTCCTCGC CCGCCTTCGTACT3’; S2-5’CTGTATATACGTGTATATATGTTGTGTGTATGTGAAT GAGCAGTGGATATCGATGAATTCGAGCTCGTTCCTCGCCCGCCTTCGTACT3’). *CDC55-GFP-NLS* and *CDC55-GFP-NES* strains were constructed by isolating the C-terminal region of *CDC55* from strains SY1808 and SY1811 (S. Yoshida) by PCR and transforming them into a W303 background using the (Forward Primer: 5’ GGGAACCGAAATG AATGAAATCG3’ Reverse Primer: 5’TCCTTTGATAGGAGTATTTGGGCGG 3’). *chk1Δ* was obtained from Euroscarf (Euroscarf, Oberursel, Germany), and PCR product using Chk1-A and Chk1-D primers were transformed into W303 strain. Standard cross methods were used to construct *rad53Δ sml1Δ* and *swe1Δ* strains using YGP24 and YGP98, respectively (D. Quintana). *pds1-38-3HA* strains were constructed by crossing with RA2815 (O. Cohen-Fix), and *pds1-5A-3HA* strains were constructed using strain LH505 (L. Holt). *PDS1-GFP* was constructed by crossing with BL123 (J. Haber).

### Cell Culture and Media

Yeast extract peptone media with glucose (YPD) was used for cell culture for western blot, flow cytometry, and serial dilution experiments. Synthetic Complete (SC) media with glucose was used for fluorescence microscopy experiments with *TUB1-GFP* strains. SC-low fluorescence media with glucose was used for fluorescence microscopy experiments with CenIV-GFP dots (*34*). Cell culture was performed at 30°C.

### Serial dilutions

Cells were grown overnight in 3mL YPD cultures at room temperature. Cells were diluted 10-fold and 5μl was spotted on YPD plates containing the indicated concentration of HU. Plates were incubated for 3 days at room temperature.

### Flow Cytometry

Cell cycle profiles were monitored by flow cytometry using propidium iodide staining as previously described (*35*). Flow cytometry analysis was performed using a BD Accuri C6 flow cytometer (BD Biosciences, San Jose, CA). 20,000 cells were analyzed per sample. Results were analyzed by FlowJo software (FlowJo LLC, Ashland, OR).

### Microscopy

For all microscopy with the exception of time lapse microscopy (Figure 6), images were obtained with a Nikon Eclipse 90i fluorescence microscope using a 60×/1.45 numerical aperture Plan Apochromatic objective lens (Nikon, Tokyo, Japan) with an Intensilight Ultra High Pressure 130-W mercury lamp (Nikon, Tokyo, Japan). Images were taken with a Clara interline charge-coupled device camera (Andor, Belfast, United Kingdom). The images were captured with NIS-Elements software (Nikon, Tokyo, Japan).

For *TUB1-GFP* time course experiments, images were captured using the DIC filter at 80ms exposure and the FITC filter for 200ms exposure (Figures 5 and 7). For CenIV-GFP dots, Z-stacks were generated with seven 0.5μM steps using the DIC filter at 80ms exposure and FITC filter at 500ms exposure. Images of flow cytometry samples using fixed cells were taken using the DIC filter at 80ms exposure time and the TxRed filter at 50ms exposure.

For time lapse microscopy, an Observer Z1 (Zeiss, Jena, Germany) microscope equipped with an automated stage and a plan-apo 63x/1.4NA oil immersion objective was used (Figure 6). Asynchronous cells were sonicated at room temperature for five seconds and transferred to 1.5% low-melting agarose pads made with SC media containing glucose. Live cell imaging was performed over 5 hours with three minutes per frame. All images were prepared using FIJI software (NIH, Bethesda, MD) (*36*). The cutoff point for long spindles was when spindle length was at its maximum length for each individual cell.

### Western blot, Co-immunoprecipitation and Phos-tag Analysis

Cells were lysed in TBT Buffer containing inhibitors with glass bead agitation as previously described (*37*). Proteins were separated using SDS–PAGE with Novex 4–20% Tris-glycine polyacrylamide gel (Invitrogen, Life Technologies, Carlsbad, CA). Western blot analysis was performed using peroxide-conjugated anti-hemaglutinin antibody at 1:250 dilution (Roche), anti-cMYC antibody 9E10 (Sigma-Aldrich, St. Louis, MO) at 1:5000 dilution, and anti-Pgk1 (Life Technologies, Carlsbad, CA) at 1:5000 as a loading control. Images were developed using a Fuji LAS 4000 Imager (GE Healthcare Life Sciences, Pittsburg, PA). Co-immunoprecipitation was performed by incubating cell lysate with with anti-MYC conjugated agarose beads for one hour at 4°C (Sigma-Aldrich, St. Louis, MO). Phos-tag analysis was performed using phosphoproteins obtained by TCA precipitation and separated with phos-tag acrylamide gels as previously described (Fujifilm Wako Pure Chemical, Osaka, Japan). (*38–40*).

### In vitro phosphatase assay

Pds1-3HA was immunoprecipitated and phosphorylated by Cdc28^Clb2^. The purification of Cdc28^Clb2^ complex was performed as previously described (*26*, *41*). Kinase reactions were performed with 1 μCi γ-[^32^P]ATP (*26*). Phosphorylated Pds1-3HA was then treated with TAP-purified PP2A^Cdc55^ complexes (*26*, *42*).

## Results

### Nuclear PP2A^Cdc55^ is involved in the replication stress response

There are four possible scenarios for PP2A^Cdc55^ function in Pds1 stabilization during replication stress (Figure 1A). In scenario 1, PP2A^Cdc55^ acts in the Intra-S checkpoint as a downstream component of Mec1 to stabilize Pds1 or to inhibit CDK. In scenario 2, PP2A^Cdc55^ acts through the Swe1 pathway to inhibit mitotic CDK activity. In scenario 3, PP2A^Cdc55^ acts in the SAC pathway to dephosphorylate APC components Cdc16 and Cdc27. Dephosphorylation of Cdc16 and Cdc27 inhibits APC^Cdc20^ activity and stabilizes Pds1. In scenario 4, PP2A^Cdc55^ is involved in a novel signaling pathway and directly regulates Pds1.

We performed serial dilution assays using a *cdc55Δ* strain on plates containing the dNTP-depleting agent hydroxyurea (HU) to test if PP2A^Cdc55^ is involved in the replication stress response *in vivo*. It confirmed a growth defect in the *cdc55Δ* strain on HU plates (Figure 1B, lane 2) (*31*). Because PP2A^Cdc55^ activity is dependent on Cdc55 localization, we tested HU sensitivity of the *cdc55-101* mutant, which excludes Cdc55 from the nucleus (Figure 1B, lane 4) (*32*). The *cdc55-101* mutant showed the same degree of HU sensitivity as *cdc55Δ* (Figure 1B, lanes 2 and 4). Similarly, *CDC55-NES* which contains a nuclear export signal exhibited the same sensitivity to HU as *cdc55Δ* (Supplemental Figure 1A, Lanes 1 and 4). *CDC55-NLS* containing nuclear localization signal showed a severe growth defect on HU, indicating that both nuclear and cytoplasmic Cdc55 are involved in replications stress response, but likely act in separate pathways (Supplemental Figure 1A, Lanes 1 and 3)

To test if PP2A^Cdc55^ acts in one of the known replication stress response pathways, we combined *cdc55-101* with mutations in the Intra-S checkpoint including *mec1Δ sml1Δ* and *rad53Δ sml1Δ* (Figures 1A Scenario 1, C-D). *MEC1* and *RAD53* are essential genes, but their lethality in deletion strains can be rescued by adding *sml1Δ* (*43*). It is known that the *mec1Δ sml1Δ* double mutant is highly sensitive to low concentration of HU at 2.5mM (Figure 1C, lane 1) (*1*). *mec1Δ sml1Δ* showed a more severe growth defect than *cdc55-101* on HU, making it unlikely that PP2A^Cdc55^ acts upstream of Mec1 (Figures 1C, lanes 1 and 4). The triple mutant *cdc55-101 mec1Δ sml1Δ* showed synthetic lethality compared to the double mutant *mec1Δ sml1Δ* (Figure 1C, lanes 3 and 5). Confirming that *sml1Δ* alone does not show HU sensitivity, the *Δsml1* single mutant did not show HU sensitivity and the *cdc55-101 sml1Δ* double mutant did not show a genetic interaction (Figure 1C, lanes 6 and 10). This enhanced growth defect of *cdc55-101 mec1Δ sml1Δ* supports a role for PP2A^Cdc55^ that is independent of the Mec1 pathway. This was further supported by evidence that the *cdc55-101 rad53Δ sml1Δ* mutant also showed synthetic lethality compared to the *rad53Δ sml1Δ* mutant (Figure 1D, lanes 2 and 4).

Mec1 activates a second effector kinase, Chk1, during MMS-dependent DNA damage response to phosphorylate and stabilize Pds1(*3*). We tested if *chk1* deletion has a genetic interaction with *cdc55-101*. *chk1Δ* cells did not show sensitivity to HU even at high concentration and there was no genetic interaction between *chk1Δ* and *cdc55-101* (Supplemental Figure 1B, lanes 1, 4 and 5). This is consistent with previous results that Chk1 is not involved in replication stress response (*3*). The M-CDK inhibitor Swe1 was recently proposed to be involved in the replication stress response in a pathway redundant to Rad53 (Figure 1A, Scenario 2) (*9*). Sensitivity to HU was tested using *swe1Δ cdc55-101* cells. A mild synthetic growth defect was observed in *swe1Δ cdc55-101* cells compared to *cdc55-101* cells at 100mM HU, suggesting that these mutations act in separate pathways (Supplemental Figure 1C, lanes 4 and 5). Furthermore, deletion of *MIH1* was synthetic lethal with *cdc55-101* on YPD plates (Supplemental Figure 1D). As Mih1 counteracts M-CDK inhibitory phosphorylation by Swe1, these results confirm a role for nuclear PP2A^Cdc55^ that is independent of Swe1-dependent CDK inhibition.

Next, we tested if PP2A^Cdc55^ plays a role in the SAC pathway (Figure 1A, Scenario 3) (*26*). The SAC-deficient *mad2Δ* strain did not show HU sensitivity, indicating that the SAC is dispensable for the replication stress response *in vivo* (Supplemental Figure 1E, lane 1). Moreover, the growth defect in *cdc55-101* was identical to the growth defect in *cdc55-101 mad2Δ*, indicating that the *cdc55-101* mutation was solely responsible for HU sensitivity in the double mutant (Supplemental Figure 1E, lanes 4 and 5). In addition, we examined the phosphorylation status of the APC component Cdc16 during HU treatment using a phos-tag reagent to better separate phosphoisoforms for immunoblotting analysis (*38*). Cdc16 was hyperphosphorylated in *cdc55-101* compared to WT when cells are untreated at time 0, as previously reported (Supplemental Figure 1F, arrow) (*26*). Cdc16 phosphorylation status was unchanged after HU treatment, suggesting that phosphorylation of this APC component is not a target of the replication stress response (Supplemental Figure 1F). Taken together, our data indicate that PP2A^Cdc55^ has a role independent of SAC during replication stress.

### Sister chromatids remained cohesed in *cdc55-101* cells during replication stress

Replication stress response results in cell cycle arrest with unsegregated sister chromatids. Because PP2A^Cdc55^ plays a role in SAC to arrest cell cycle in metaphase, we tested if *cdc55-101* mutant affects cell cycle and chromosome segregation during replication stress. Wild type (WT) and *cdc55-101* cells were synchronized in G1 by alpha-factor and released into media containing 100mM HU (Figure 2A-B). Cell cycle progression was slightly accelerated in the *cdc55-101* mutant compared to WT starting at 120 minutes (Figure 2A). We examined sister chromatid segregation in propidium iodide-stained cells by fluorescence microscopy (Figure 2B). While neither WT nor *cdc55-101* cells showed significant chromosome segregation during the course of HU treatment, a “stretched” chromatid pattern was observed in *cdc55-101* cells (Figure 2B, red bar). The stretched staining pattern was found in large-budded cells where connected staining extended to both cell bodies (Figure 2B, right). At 180 minutes, 39% of *cdc55-101* cells exhibited the stretched phenotype, compared to 18% in WT (Figure 2B).

**Figure 2.**
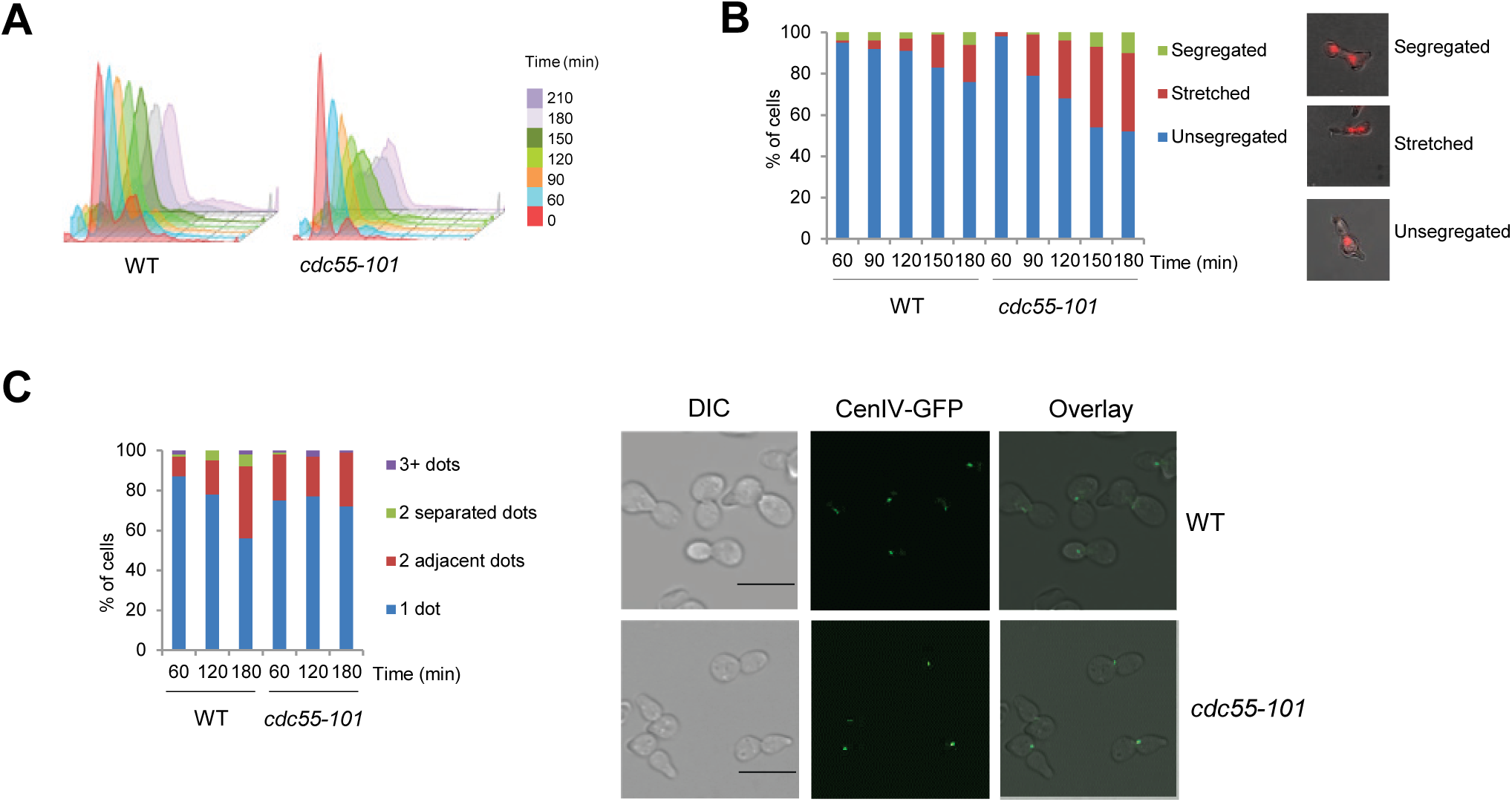
Sister chromatids remained cohesed in *cdc55-101* cells at 100mM HU. (A) WT or *cdc55-101* cells were arrested in G1 by alpha factor and released into YPD containing 100mM HU. Samples were taken at the indicated time points. Cell cycle profile was determined by flow cytometry. (B) Sister chromatid segregation status of samples obtained in (A) was analyzed by fluorescence microscopy. Propidium iodide-stained cells were counted and categorized based on status as; Green:fully segregated chromosomes, Red:stretched chromosomes and Blue: unsegregated chromosomes. Representative images from t=120 are shown (Right). (C) *CDC55 promURA::tetR::GFP cenIV::tetOx448* (WT) or *cdc55-101 promURA::tetR::GFP cenIV::tetOx448* (*cdc55-101)* cells were arrested in G1 by alpha factor and released into 100mM HU. GFP dots were counted per cell at the indicated time points. A total of 100 cells were counted and the percentage of the cells for each category is shown. Blue indicates cells with1 dot, Red indicates cells with 2 adjacent dots, Green indicates cells with 2 separated dots and Purple indicates cells with 3 or more dots. Representative images at 120 minutes after the release are shown on the right. Scale bar represents 10µM.

Next, we tested if the stretched chromosome pattern was due to partial chromosome segregation. A TetO-TetR system was used to visualize the centromere at Cen IV (*44*, *45*). GFP dots on Cen IV were counted at the indicated times during HU treatment after G1 release. There was no significant difference in centromere separation between WT and *cdc55-101* cells which supports the idea that chromosomes are not prematurely segregated in the *cdc55-101* mutant (Figure 2C).

### Excluding PP2A^Cdc55^ from the nucleus results in Pds1 hyperphosphorylation

Pds1 is a main target in replication stress response. Since Pds1 is a CDK substrate, and PP2A^Cdc55^ counteracts CDK phosphorylation events, we tested if PP2A^Cdc55^ regulates Pds1 phosphorylation status (*17*, *22*). A time course experiment was performed as described in Figure 2A to examine Pds1 phosphorylation status (Figure 3A). Pds1 protein migrated as two distinct bands, with the upper band presumably representing the hyperphosphorylated form, as previously reported (*17*). The *cdc55-101* mutant showed a higher ratio of hyperphosphorylated to hypophosphorylated Pds1 protein compared to WT (Figure 3A). At a lower concentration of HU (50mM), which limits Intra-S checkpoint activation, Pds1 hyperphosphorylation was observed at time points 60, 90, and 120 minutes after HU addition in *cdc55-101* cells which was less pronounced than at the higher HU concentration (Figure 3B, Right). Therefore, Pds1 hyperphosphorylation was enhanced in *cdc55-101* mutant in a HU dose-dependent manner. It indicates that PP2A^Cdc55^ may limit Pds1 phosphorylation when cells are treated with high concentration of HU.

**Figure 3.**
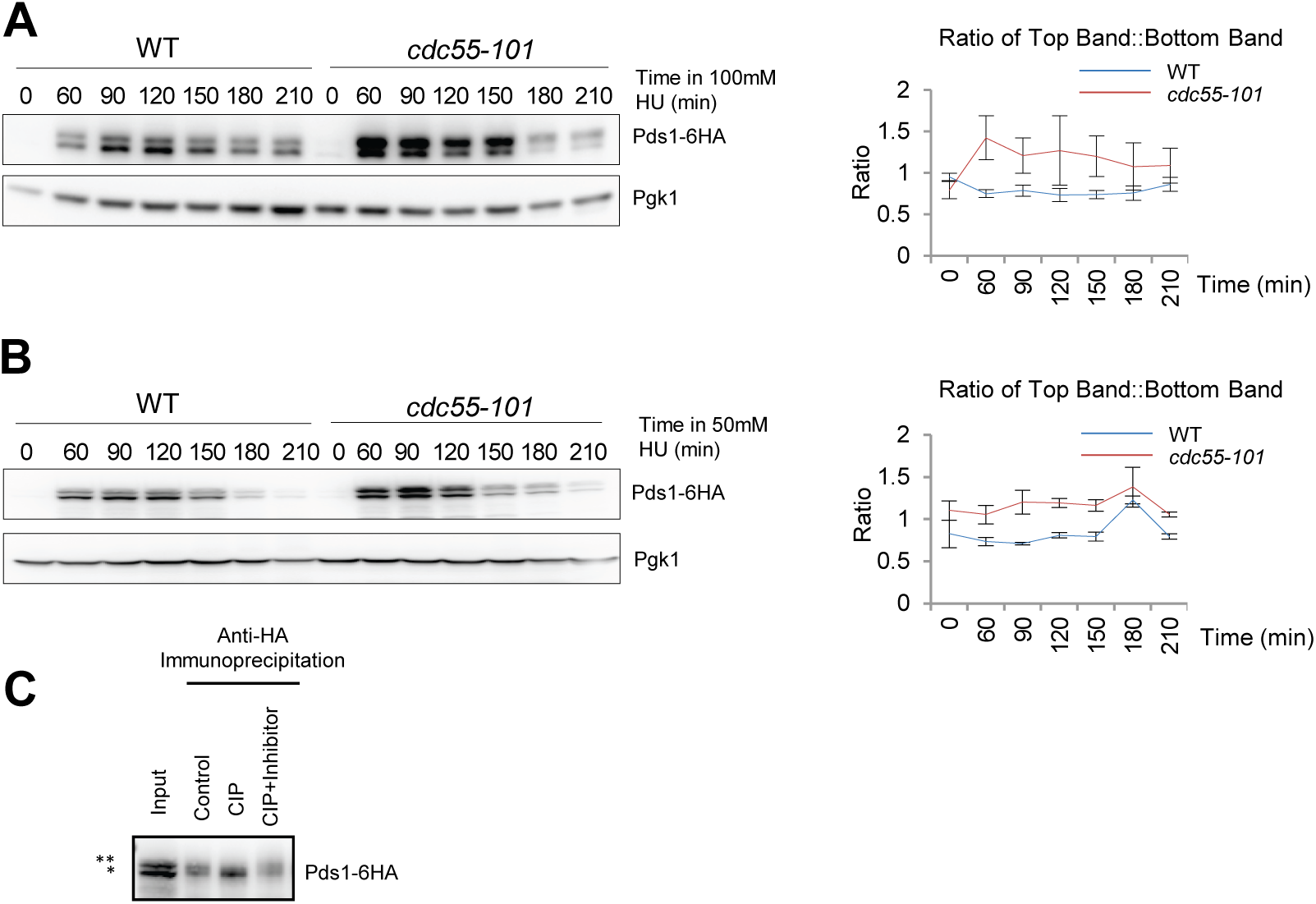
Pds1 is hyperphosphorylated in the *cdc55-101* mutant in a HU-dose-dependent manner. (A) *PDS1-HA (WT)* or *PDS1-HA cdc55-101 (cdc55-101)* cells were arrested in G1 phase by alpha factor, then released into YPD media containing 100mM HU. Alpha factor was added to the cultures again at 80 minutes after release. Cells were collected for Western blot at the indicated times. Pgk1 was used as a loading control. Pds1 upper and lower bands were quantified. A ratio of top band to bottom band were calculated from three independent experiments. Blue= WT; Red= *cdc55-101*. Error bars represent the SEM (right). (B) A time course was performed as described in A, except using 50mM HU. (C) *cdc55Δ swe1Δ PDS1-6HA* cells were grown in YPD, arrested in G1 with alpha factor, and released into YPD for 45 minutes. Cells were lysed and anti-HA immunoprecipitation was performed. IP samples were incubated at 37°C for 15 minutes without CIP, in the presence of CIP, and in the presence of CIP and phosphatase inhibitors. * represents unphosphorylated protein. ** represents phosphoprotein. Protein samples were analyzed by western blotting.

We also confirmed that the Pds1 upper band is hyperphosphorylated Pds1. Pds1 protein was immunoprecipitated from *cdc55Δ* cell lysate and treated with calf intestinal phosphatase (CIP) in the presence or absence of a phosphatase inhibitor (Figure 3C). The top Pds1 band (shown with **) was absent in CIP-treated cells, but present when the phosphatase inhibitor was added (Figure 3C). Thus, we concluded that the top band represented hyperphosphorylated Pds1. To test if Pds1 hyperphosphorylation was specifically due to PP2A^Cdc55^, we deleted the only other PP2A B-regulatory subunit, Rts1 (*21*). In *rts1Δ* cells, there were no changes in Pds1 phosphorylation patterns between WT and *rts1Δ* cells (Supplemental Figure 2). Taken together, these findings demonstrate a specific role for PP2A^Cdc55^ in regulating Pds1 phosphorylation during replication stress.

### PP2A^Cdc55^ directly dephosphorylates Pds1 *in vitro*

Next, we tested if purified PP2A^Cdc55^ dephosphorylates Pds1 *in vitro*. First, Pds1-3HA was immunoprecipitated from a WT strain and incubated with purified Clb2-Cdc28 complex *in vitro*, resulting in Pds1 phosphorylation (Figure 4A, lane 3). This Pds1 phosphorylation was reduced when the Pds1-5A variant protein, which contains S/T→A mutations at all five CDK consensus sites (T27A, S71A, S277A, S292A, T304A) (Figure 4A, lane 2)(*46*). The phosphorylated wild type Pds1-3HA from lane 3 was then incubated with purified PP2A^Cdc55^. The Pds1 was fully dephosphorylated between 30-60 minutes after the incubation (Figure 4B). When the phosphorylated Pds1 was incubated without purified PP2ACdc55, it restored the phosphorylation even after 60 minutes. This result confirms that PP2A^Cdc55^ directly dephosphorylates Pds1 *in vitro*.

**Figure 4.**
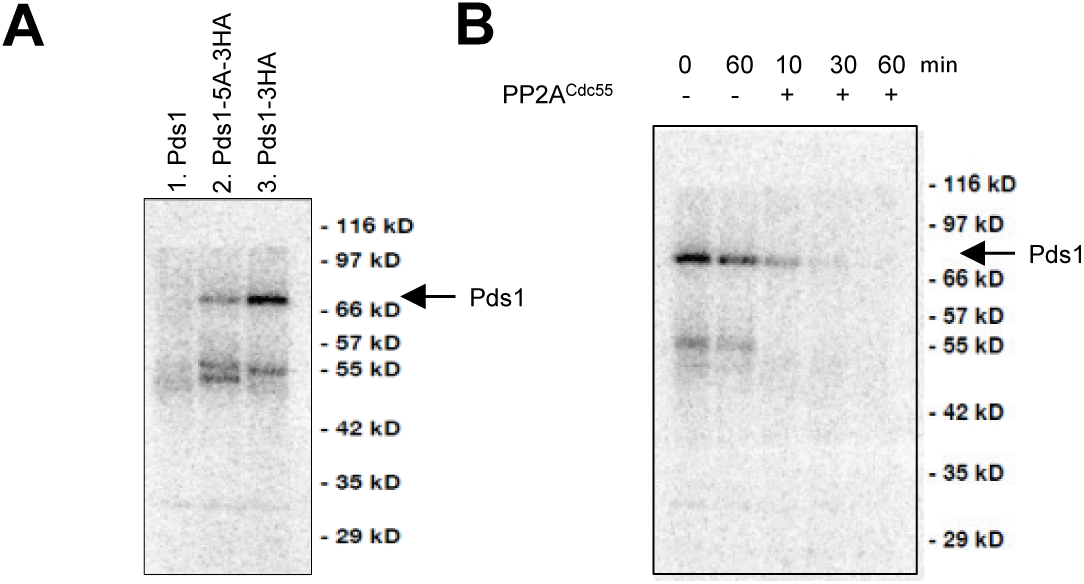
PP2A^Cdc55^ dephosphorylates Pds1. (A) Purified Pds1-3HA protein was incubated with Clb2^Cdc28^ complex and γ-32. Pds1-5A-3HA contains mutations at CDK consensus sites. Non-tagged strain (Pds1) was used as a negative control. Samples were subjected to SDS-PAGE to detect γ-32P. (B) TAP-purified PP2A^Cdc55^ was added to the phosphorylated Pds1-3HA extract in (A, lane 3). Samples were collected 10, 30 or 60 minutes after incubation at room temperature and subjected to SDS-PAGE to detect γ-32P. The phosphorylated Pds1-3HA was incubated without PP2A^Cdc55^ and was incubated for 60min as a control.

### PP2A^Cdc55^ nuclear exclusion is associated with premature spindle elongation

We next determined how Pds1 phosphorylation status affects spindle elongation during anaphase. Pds1 C-terminal phosphorylation is necessary for the physical interaction with Esp1 which plays a role in Pds1 nuclear localization (*17*, *18*). We did not observe premature sister chromatid separation in *cdc55-101* cells, however there was stretched chromosome phenotype *cdc55-101* cells under HU treatment (Figure 2B-C). We hypothesized that PP2A^Cdc55^ may affect spindle dynamics. *TUB1-GFP* was used to visualize tubulin in WT and *cdc55-101* cells in the presence of 50mM HU. The *cdc55-101* cells showed accelerated long spindle formation at 90 minutes, while WT cells did not form long spindles until 120 minutes (Supplemental Figure 3A). The same samples were subjected to western blotting to analyze the Pds1 level and phosphorylation status. Pds1 was hyperphosphorylated between 90 to 120 minutes which coincides with the accelerated spindle formation (Figure 3B and Supplemental Figure 3A).

In order to examine fully elongated spindles, Pds1 phosphorylation and spindle elongation status were monitored without HU. After 60 minutes from G1 release, *cdc55-101* showed 31%±7.3 of cells exhibiting long spindles compared to 13.3%±1.9 in WT (Figure 5A). In addition, *cdc55-101* cells showed a stronger Pds1 hyperphosphorylation band compared to WT at 45 and 60 minutes following G1 release in the same samples (Figure 5B) The timing of spindle elongation in *cdc55-101* cells corresponded to time points where Pds1 was hyperphosphorylated (Figure 5A-B). Therefore spindle elongation was accelerated in the *cdc55*-*101* mutant compared to WT in unperturbed cell cycle.

**Figure 5.**
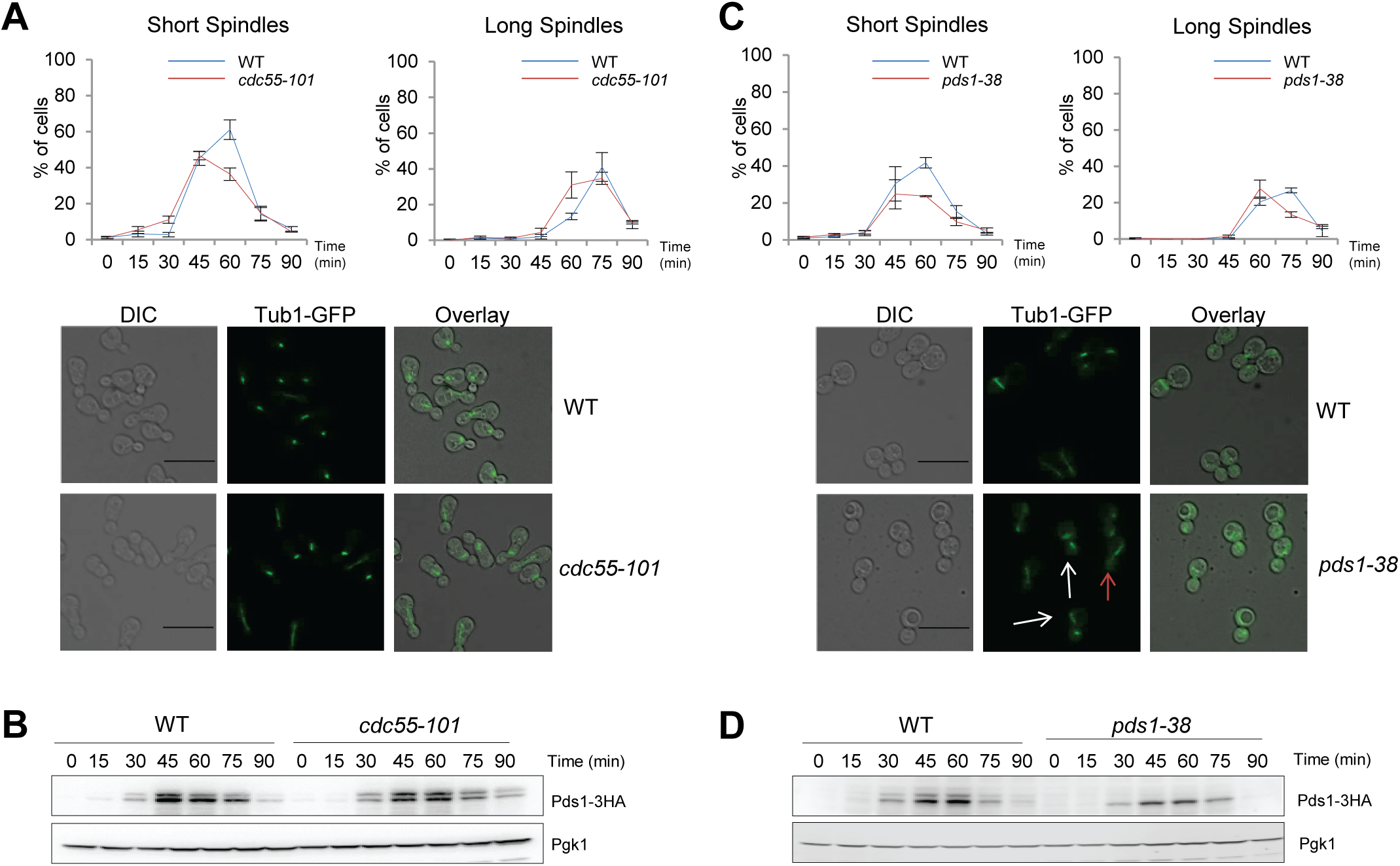
*cdc55-101* and *pds1-38* cells show altered spindle dynamics compared to WT. (A) *TUB1-GFP* (WT) or *TUB1-GFP cdc55-101* (*cdc55-101*) cells were arrested in G1 by alpha factor and released into synthetic complete media. Alpha factor was added again at 55 minutes after release. The number of cells exhibiting shorts spindles (left) and long spindles (right) were counted for a total of 100 cells per time point. Graphs show the average of three independent experiments. Error bars represent SEM. Blue=WT; Red=*cdc55-101*. Representative images from t=60 min are shown (bottom). Scale bar represents 10µM. (B) The same samples from (A) were subjected to western blotting to observe Pds1. Pgk1 was used as a loading control. (C) *TUB1-GFP* (WT) or *TUB1-GFP pds1-38 (pds1-38)* cells were treated and analyzed as described in (A). Graphs show the average of three independent experiments. Blue=WT; Red=*pds1-38*. Representative images from t=75 min are shown (bottom). (D) The same samples from C were subjected to western blotting to observe Pds1. Pgk1 was used as a loading control.

In order to examine spindle formation dynamics in live cells, we observed *TUB1-GFP* in WT and *cdc55-101* cells using time-lapse microscopy. The amount of time spanning from a single Tub1-GFP dot (red arrow) to a fully elongated spindle (labeled as long spindle) was measured in individual cell (Figure 6A-B). The average time to form long spindles was 79 minutes in WT cells. The *cdc55-101* cells reached full spindle elongation at 68 minutes which was 11 minutes faster than WT on average (Figure 6D). Taken together, these data show that *cdc55-101* mutant accelerates spindle elongation.

### Pds1 C-terminal phosphorylation mutant shows fragile mitotic spindles

We next sought to determine which Pds1 phosphorylation sites affect spindle dynamics. Pds1 C-terminal phosphorylation facilitates a physical interaction with Esp1 (*17*). Because Esp1 is localized to the spindles, we hypothesized that Pds1 C-terminal phosphorylation is necessary for spindle elongation. We examined spindle elongation in WT and *pds1-38* cells, which contain three S/T→A mutations at the C-terminus CDK phosphorylation sites. The number of cells with short spindles in *pds1-38* cells was reduced compared to WT (Figure 5C). At 60 minutes, 23%±0.3 of *pds1-38* cells had short spindles, compared to 41%±2.8 in WT cells (Figure 5C, Left). At 75 minutes, both strains formed long spindles; 13%±1.5 of *pds1-38* versus 27%±1.3 in WT (Figure 5C, Right). Long spindles in *pds1-38* cells were structurally aberrant from spindles in WT cells. The *pds1-38* strain exhibited asymmetric long spindles (Figure 5C, red arrow), which were more likely to separate unevenly between the mother and daughter cells (Figure 5C, white arrows). These results indicate that both short and long spindles were unstable in *pds1-38* cells, which was the opposite of what was seen in *cdc55-101* cells. We also confirmed that the hyperphosphorylated form of Pds1 was fully abolished in the *pds1-38* mutant (Figure 5D). From these experiments, we conclude that Pds1 C-terminal phosphorylation status regulates spindle elongation and morphology.

Next, we monitored spindles in *pds1-38* cells using time-lapse microscopy (Figure 6C). There was no significant difference in the length of time from the appearance of the initial Tub1-GFP (first image with red arrow) single dot to a fully elongated spindle (labeled as Long Spindle) (Figure 6D). Consistent with findings from time course experiments in Figure 5C, *pds1*-*38* cells had an irregular spindle shape when elongated (Figure 6C, blue arrow). Short spindles in *pds1-38* cells also moved between the mother and daughter cell bodies before elongating (Figure 6C, yellow arrows). This shuttling movement was seen in 41% of *pds1-38* cells compared to 23% in WT cells.

**Figure 6.**
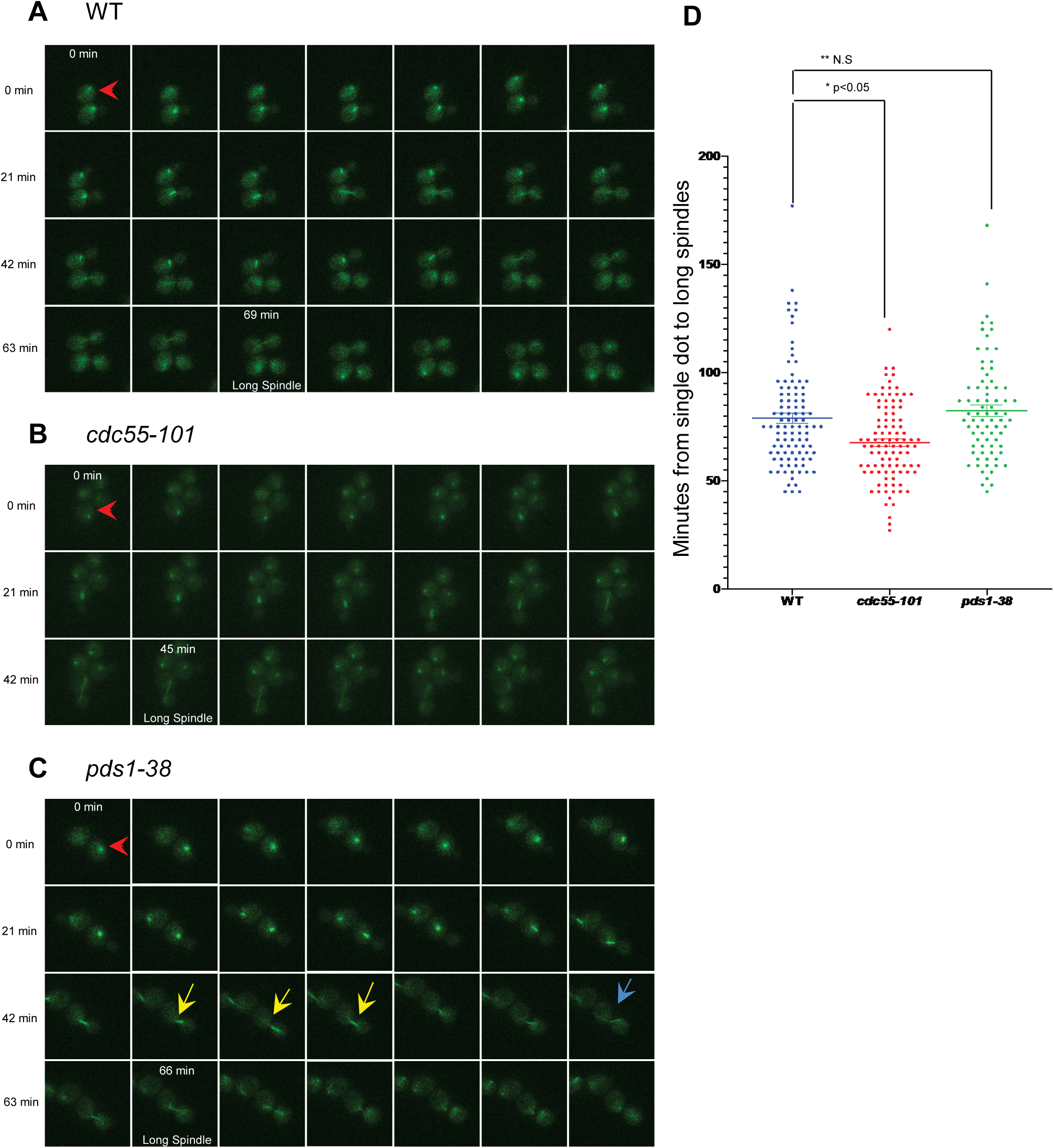
*pds1-38* and *cdc55-101* show altered spindle elongation rates and morphology. *TUB1-GFP CDC55* (A), *TUB1-GFP cdc55-101* (B), and *TUB1-GFP pds1-38* (C) were observed by time-lapse fluorescence microscopy. Cell images were taken every three minutes. Red arrows indicate the time point when cells showed spindle formation. Fully elongated spindles are labeled as “Long Spindle”. Yellow arrows show spindle shuttling movement. Blue arrow shows irregular spindle shape. (D) Time from initial presence of GFP dot to fully elongated spindle was monitored per cell. WT, *cdc55-101*, and *pds1-38* strains were used (n=100, 100, 78, respectively). *\* p<.05 **Not significant*

To confirm if Pds1 C-terminal phosphorylation sites are targeted by PP2A^Cdc55^, we used a *pds1-38 cdc55-101* double mutant strain to examine Pds1 phosphorylation status and spindle morphology. We found that spindle elongation rates in *pds1-38 cdc55-101* double mutant cells were similar to that in the *pds1-38* single mutant (Figure 7A, left and middle). The *cdc55-101 pds1-38* double mutant also exhibited the irregular spindle structure that was present in the *pds1*-*38* single mutant (Figure 7A, white arrow). There was no Pds1 phosphorylation in either the *pds1-38* single mutant or the *pds1-38 cdc55-101* double mutant suggesting for a role of Pds1 in regulating spindle dynamics downstream of PP2A^Cdc55^ (Figure 7B). Under replication stress, most of the phosphorylated Pds1 band was also eliminated in the *cdc55-101 pds1-38* double mutant (Supplemental Figure 4A). We further used *pds1-5A cdc55-101* cells in which both N- and C-terminal CDK phosphorylation sites are mutated. All of the *pds1-38* mutation sites are included in *pds1-5A*, in addition to two N-terminal S/T→A mutations at T27 and S71 (*46*). The phosphorylated form of Pds1 was completely abolished in *pds1-5A* confirming our idea PP2A^Cdc55^ targets Pds1 at CDK sites (Supplemental Figure 4B). Taken together, these results demonstrate that PP2A^Cdc55^-dependent Pds1 dephosphorylation is limited to the five CDK consensus sites.

**Figure 7.**
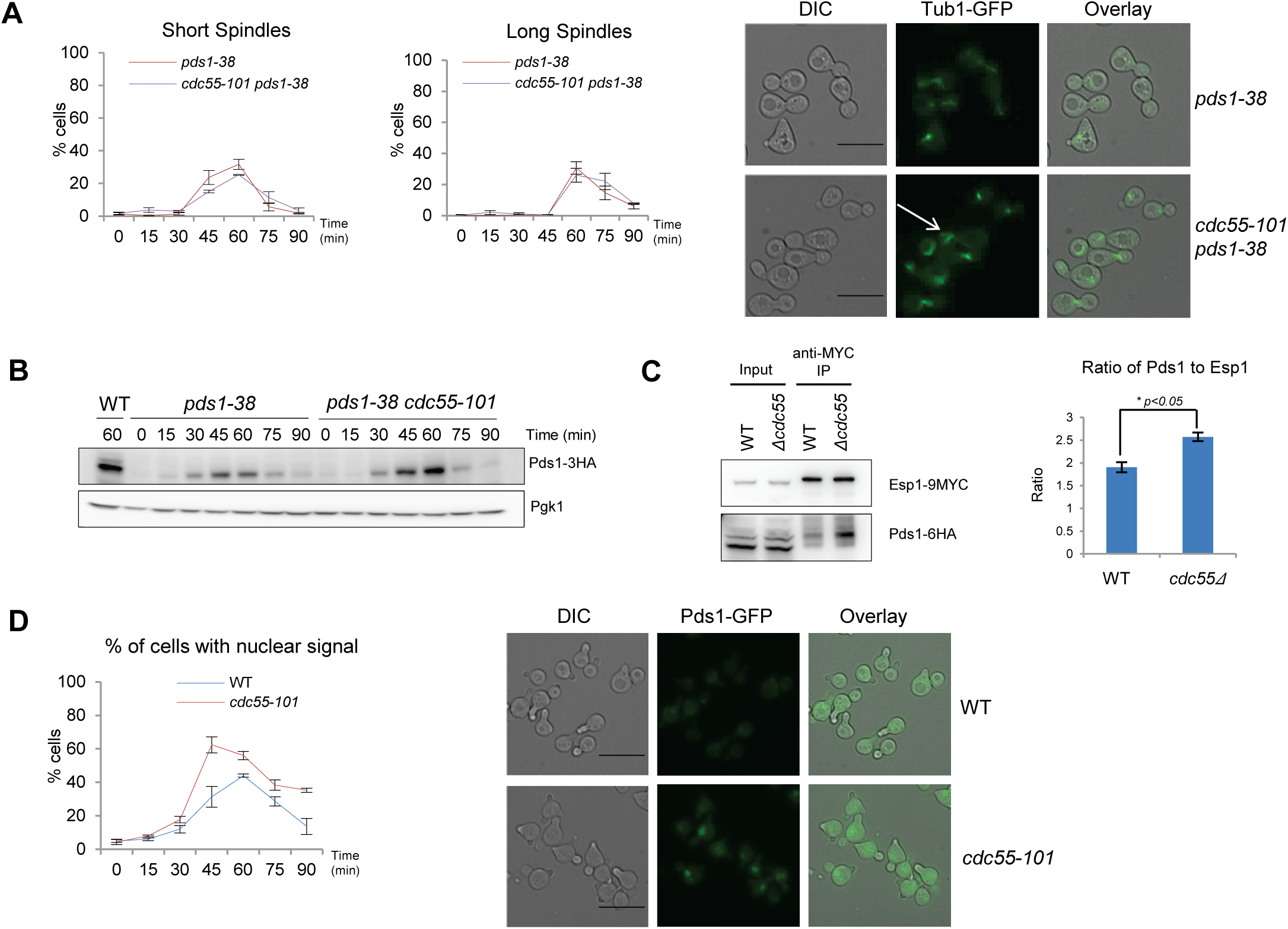
Pds1 phospho-regulation affects spindle morphology, Pds1-Esp1 association, and Pds1 localization. (A) *pds1-38 TUB1-GFP (pds1-38)* and *cdc55-101 pds1-38 TUB1-GFP (cdc55-101 pds1-38)* cells were examined by fluorescence microscopy in a time course as described in Figure 5A. Images were taken at the indicated time points. The number of cells exhibiting short spindles (Left) and long spindles (Middle) were counted. Graphs represent the average of 3 independent experiments. Error bars represent SEM. Red= *pds1-38*; Purple= *pds1-38 cdc55-101*. Representative images from t=75 are shown (Right). Scale bar represents 10µM. (B) Pds1-3HA levels and phosphorylation status in *pds1-38-3HA* and *cdc55-101 pds1-38-3HA* cells were examined by western blot analysis during a time course as described in Figure 5B. WT Pds1-3HA from a synchronized culture is shown as a control (Lane 1). (C) (Left) *PDS1-6HA ESP1-9MYC* (WT) and *PDS1-6HA ESP1-9MYC cdc55Δ (cdc55Δ)* cells were grown asynchronously. Immunoprecipitation was performed with anti-MYC agarose beads. Esp1-MYC and Pds1-HA levels were detected by western blot. (Right) Ratio of HA signal intensity to MYC signal intensity is shown. Chart represents average of three independent experiments. Error bars represent SEM. *\*p<.05* (Student’s *t* test) (D) *PDS1-GFP* (WT) and *PDS1-GFP cdc55-101* (*cdc55-101*) cells were grown in low fluorescence media, arrested by alpha-factor. Cell cycle arrest was released and alpha-factor was added again at 55 minutes after release. (Left) The graph shows percentage of the cells with Pds1 nuclear signal. An average percentage from three independent experiments is shown. Error bars represent SEM. Blue=WT, Red=*cdc55-101*. (Right) Representative images from t=45 are shown.

Since phosphorylated Pds1 was reported to bind Esp1, we tested if the Pds1-Esp1 physical interaction is affected in *cdc55Δ* mutant (*17*). A co-IP experiment was performed using asynchronous WT and *cdc55Δ* cells. Esp1-9MYC was immunoprecipitated and the associated Pds1 protein was monitored. The protein ratio of Pds1 to Esp1 in the IP was used to determine if the Pds1-Esp1 interaction is altered in the absence of Cdc55. Our findings show that the Pds1-Esp1 physical interaction was enhanced in the *cdc55Δ* strain (Figure 7C). Taken together, these results implicate a role for PP2A^Cdc55^ in disrupting the Pds1-Esp1 interaction through Pds1 dephosphorylation.

To test if Cdc55 affects Pds1 localization, we examined *PDS1-GFP* localization in WT and *cdc55-101* cells under fluorescent microscope. Pds1 was localized to the nucleus earlier in *cdc55-101* compared to WT cells (Figure 7D). At 45 minutes after G1 release, 62%±4.8 of *cdc55-101* cells showed Pds1 nuclear localization, compared to 31%±6.2 of WT cells (Figure 7D). At the 45-minute time point, Pds1 is more hyperphosphorylated in *cdc55-101* compared to WT, suggesting that there is a correlation between Pds1 hyperphosphorylation and nuclear localization (Figure 5B). We attempted to generate a *pds1Δ cdc55-101* strain, but double mutants were synthetic lethal, indicating that additional genetic interactions between *PDS1* and *CDC55* are present (Supplemental Figure 4C). Synthetic lethality in *pds1Δ cdc55-101* is consistent with an earlier finding that *pds1Δ cdc55Δ* double mutants are inviable (*28*).

## Discussion

### PP2A^Cdc55^ acts in a novel replication stress response pathway

Previous reports have focused on the role of PP2A^Cdc55^ in nocodazole treated cells when the spindle is disrupted (*34*). Pds1 is prematurely degraded by APC^Cdc20^ in *cdc55Δ* cells resulting in precocious sister chromatid separation in nocodazole treated cells. In this case, PP2A^Cdc55^ targets and inhibits APC^Cdc20^. This pathway relies on Esp1’s protease function to cleave Scc1. In the presence of DNA damage in *cdc13-1* cells, premature chromatid separation was independent from Pds1 degradation. However, the molecular mechanism is unclear (*28*). In this study, we used HU to activate the Intra-S checkpoint and studied how PP2A^Cdc55^ is involved in the checkpoint response. We found that PP2A^Cdc55^ directly dephosphorylates Pds1 to inhibit mitotic spindle elongation but does not affect cohesin cleavage. PP2A^Cdc55^ dependent Pds1 dephosphorylation releases Esp1 from Pds1 which may inhibit mitotic spindle formation during replication stress. We speculate that the free Esp1 is not able to cleave Scc1 because it is localized in the cytoplasm.

PP2A^Cdc55^ has distinct functions depending on its localization. Our results showed that both nuclear and cytoplasmic PP2A^Cdc55^ activities are necessary for maintaining genomic integrity during replication stress. We showed that nuclear PP2A^Cdc55^ prevents premature spindle elongation directly through Pds1, and not through Mec1-, Swe1 or Mad2-pathways. It is known that the cytoplasmic PP2A^Cdc55^ inhibits Swe1 activating M-CDK activity during unperturbed cell cycle. It has been shown that Swe1 is stabilized in response to HU, but not to nocodazole (*31*). We propose a model that replication stress activates PP2A^Cdc55^ both in nucleus and cytoplasm which contributes to the mitotic block with distinct targets. We attempted to examine Cdc55 localization during replication stress, but there were no obvious changes detected (data not shown).

### PP2A^Cdc55^ disrupts the Pds1-Esp1 interaction

PP2A^Cdc55^ regulates Pds1 phosphorylation status both in unperturbed and replication stress conditions specifically at CDK consensus sites necessary for its Esp1 interaction. Recent studies showed that the Pds1 C-terminal segment (residues 258-373) is required for the Esp1 interaction, which is consistent with the idea that phosphorylation at the C-terminal sites located at S277, S292, T304 are required for Esp1 binding (*47*). They propose a model that S277 should have favorable interactions with Esp1 due to a possible conformational change induced by S292 and T304 phosphorylation. When Pds1 and Esp1 are bound, the Pds1 C-terminal region is located in the Esp1 active site (*47*). While this explains how Pds1 may block Esp1 protease activity, it remains unclear if the interaction site affects Esp1 function in spindle elongation. It was previously proposed that the Pds1-Esp1 interaction causes a conformational change in Esp1 that is a prerequisite for Esp1 proteolytic activity, although continued Pds1 binding would prevent Esp1 from interacting with a cleavage substrate (*48*). The Esp1 C-terminus, which contains its catalytic domain, is necessary for spindle interaction and thus it is possible that an Esp1 conformational change is also a prerequisite for spindle elongation (*18*, *19*).

### Pds1 phosphorylation status regulates spindle elongation and spindle structure

The process of spindle elongation is dependent on Clb1/2-CDK (Cdc28) activity (*49*). Mitotic spindle elongation requires Clb2-Cdc28 mediated phosphorylation of the functionally redundant kinesin motor proteins Kip1 and Cin8 (*50*). Kip1 and Cin8 interact with the spindle to provide outwardly-directed force and overexpression of Cin8 results in premature spindle elongation (*51*, *52*). Both Kip1 and Cin8 contain the Esp1 core cleavage motif (D/E)XXR and show decreasing protein levels in mitosis, which raises the possibility that Esp1 cleaves them to promote their degradation (*53*). In our proposed scenario, Pds1 hyperphosphorylation enhances the Pds1-Esp1 interaction, which would keep the Esp1 active site occupied by Pds1. Without Esp1 protease activity, Kip1 and Cin8 levels should be elevated, resulting in accelerated spindle elongation.

The fragile spindles in *pds1-38* may be a result of inefficient Esp1 recruitment to the SPBs. A genetic screen using temperature-sensitive *esp1-1* showed a synthetic growth defect with overexpression of microtubule minus-end motor Kar3 and its binding partner Cik1 (*54*). Kar3 localizes to the SPB and generates inward force (*51*). It is possible that Esp1 presence at the SPB inhibits Kar3 activity to prevent spindle collapse. Therefore, excessive inwardly-directed force due to Kar3 activity may cause the observed spindle fragility in *pds1-38* cells.

The Pds1 N-terminal CDK sites (T27, S71) have unknown functions, although a potential NLS sequence is present at S71 (*55*). Premature Pds1 nuclear localization in *cdc55-101* may be a result of altered phosphorylation at S71. If hyperphosphorylated Pds1 is in a complex with Esp1, premature Pds1 localization to the nucleus may explain accelerated spindle elongation in *cdc55-101*.

### Additional genetic interactions between *CDC55* and *PDS1* are revealed

It was recently shown that PP2A^Cdc55^ dephosphorylates Esp1 at CDK consensus sites (*42*). They proposed a model in which Pds1 destruction and phospho-regulation of Esp1 have redundant functions for Esp1 activation in mitosis. Combining the phospho-mimetic mutations *esp1-3D* with *pds1Δ* resulted in synthetic lethality (*42*). Here, we found that *pds1Δ cdc55-101* is synthetic lethal, which could be due to Esp1 hyperphosphorylation in *cdc55-101* in the absence of Pds1 resulting in full activation of Esp1. While Esp1 nuclear localization would be inefficient in *pds1Δ,* this scenario would not be fully excluded (*17*). Nuclear Esp1 would be activated in the absence of both Pds1 and nuclear PP2A^Cdc55^. Synthetic lethality in *pds1Δ cdc55-101* double mutant is most likely due to uncontrolled nuclear Esp1 activity resulting in both premature Scc1 cleavage and spindle elongation.

## Conclusion

In summary, our findings show that nuclear PP2A^Cdc55^ has a novel role in regulating spindle dynamics by dephosphorylating Pds1. We propose a model where PP2A^Cdc55^ disrupts the Pds1-Esp1 interaction to inhibit spindle elongation. Pds1 dephosphorylation by PP2A^Cdc55^ has a prominent role in the replication stress response that is independent of known checkpoint mechanisms. These findings may have implications in humans as mammalian PP2A is a tumor suppressor, where mutations in human PP2A have been shown to be associated with solid tumors as well as various types of leukemia (*56–60*). The PP2A regulatory subunit B55 is the mammalian homologue of Cdc55 and human PP2A-B55δ dephosphorylates hSecurin, the human homologue of Pds1, to stabilize it (*61*, *62*). Our work here suggests it is of interest to study if hSecurin dephosphorylation by PP2A affects spindle behavior or has a role in the replication stress response.

## Acknowledgement

We gratefully acknowledge Orna Cohen-Fix for the *pds1-38-HA* strain, David Quintana for *rad53Δ sml1Δ, swe1Δ,* and *pds1Δ* strains, Satoshi Yoshida for *CDC55-MYC*, *cdc55-101-MYC*, *CDC55-GFP-NES* and *CDC55-GFP-NLS* strains, James Haber for the *PDS1-GFP* strain, and Liam Holt for *pds1-5A-3HA* strain.

A.E.I was supported by NIH 5SC1GM21242.

