## Supplementary material for "PP2A^Cdc55^ dephosphorylates Pds1 to inhibit spindle elongation": Supplmental

### Slide 1
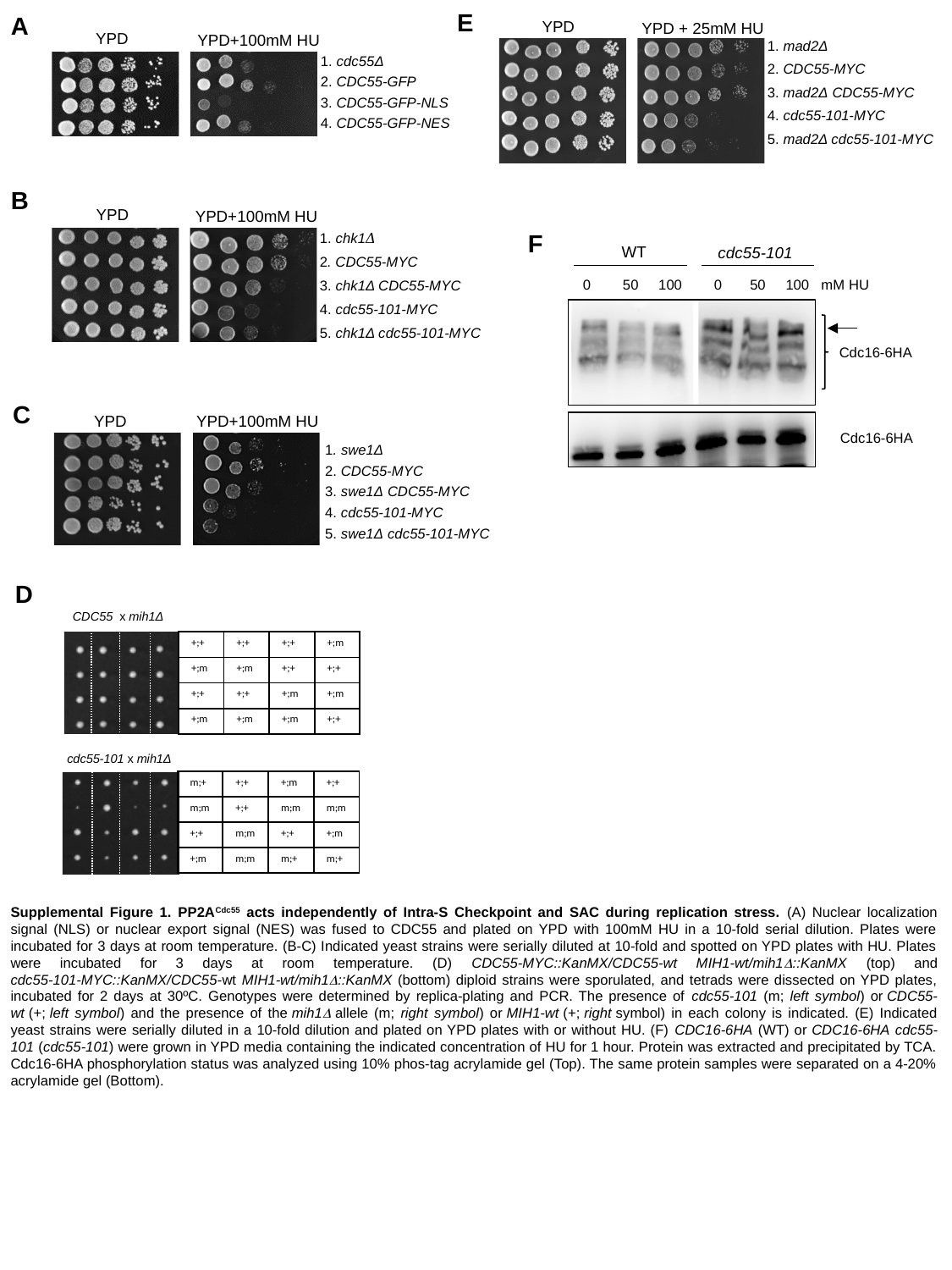

A
YPD
YPD+100mM HU
1. cdc55Δ
2. CDC55-GFP
3. CDC55-GFP-NLS
4. CDC55-GFP-NES
E
YPD
YPD + 25mM HU
1. mad2Δ
2. CDC55-MYC
3. mad2Δ CDC55-MYC
4. cdc55-101-MYC
5. mad2Δ cdc55-101-MYC
B
YPD
YPD+100mM HU
1. chk1𝛥
2. CDC55-MYC
3. chk1Δ CDC55-MYC
4. cdc55-101-MYC
5. chk1Δ cdc55-101-MYC
F
WT
cdc55-101
0 50 100 0 50 100 mM HU
Cdc16-6HA
Cdc16-6HA
C
YPD
YPD+100mM HU
1. swe1Δ
2. CDC55-MYC
3. swe1Δ CDC55-MYC
4. cdc55-101-MYC
5. swe1Δ cdc55-101-MYC
D
CDC55 x mih1Δ
| +;+ | +;+ | +;+ | +;m |
| --- | --- | --- | --- |
| +;m | +;m | +;+ | +;+ |
| +;+ | +;+ | +;m | +;m |
| +;m | +;m | +;m | +;+ |
cdc55-101 x mih1Δ
| m;+ | +;+ | +;m | +;+ |
| --- | --- | --- | --- |
| m;m | +;+ | m;m | m;m |
| +;+ | m;m | +;+ | +;m |
| +;m | m;m | m;+ | m;+ |
Supplemental Figure 1. PP2A^Cdc55^ acts independently of Intra-S Checkpoint and SAC during replication stress. (A) Nuclear localization signal (NLS) or nuclear export signal (NES) was fused to CDC55 and plated on YPD with 100mM HU in a 10-fold serial dilution. Plates were incubated for 3 days at room temperature. (B-C) Indicated yeast strains were serially diluted at 10-fold and spotted on YPD plates with HU. Plates were incubated for 3 days at room temperature. (D) CDC55-MYC::KanMX/CDC55-wt MIH1-wt/mih1Δ::KanMX (top) and cdc55-101-MYC::KanMX/CDC55-wt MIH1-wt/mih1Δ::KanMX (bottom) diploid strains were sporulated, and tetrads were dissected on YPD plates, incubated for 2 days at 30ºC. Genotypes were determined by replica-plating and PCR. The presence of cdc55-101 (m; left symbol) or CDC55-wt (+; left symbol) and the presence of the mih1Δ allele (m; right symbol) or MIH1-wt (+; right symbol) in each colony is indicated. (E) Indicated yeast strains were serially diluted in a 10-fold dilution and plated on YPD plates with or without HU. (F) CDC16-6HA (WT) or CDC16-6HA cdc55-101 (cdc55-101) were grown in YPD media containing the indicated concentration of HU for 1 hour. Protein was extracted and precipitated by TCA. Cdc16-6HA phosphorylation status was analyzed using 10% phos-tag acrylamide gel (Top). The same protein samples were separated on a 4-20% acrylamide gel (Bottom).

### Slide 2
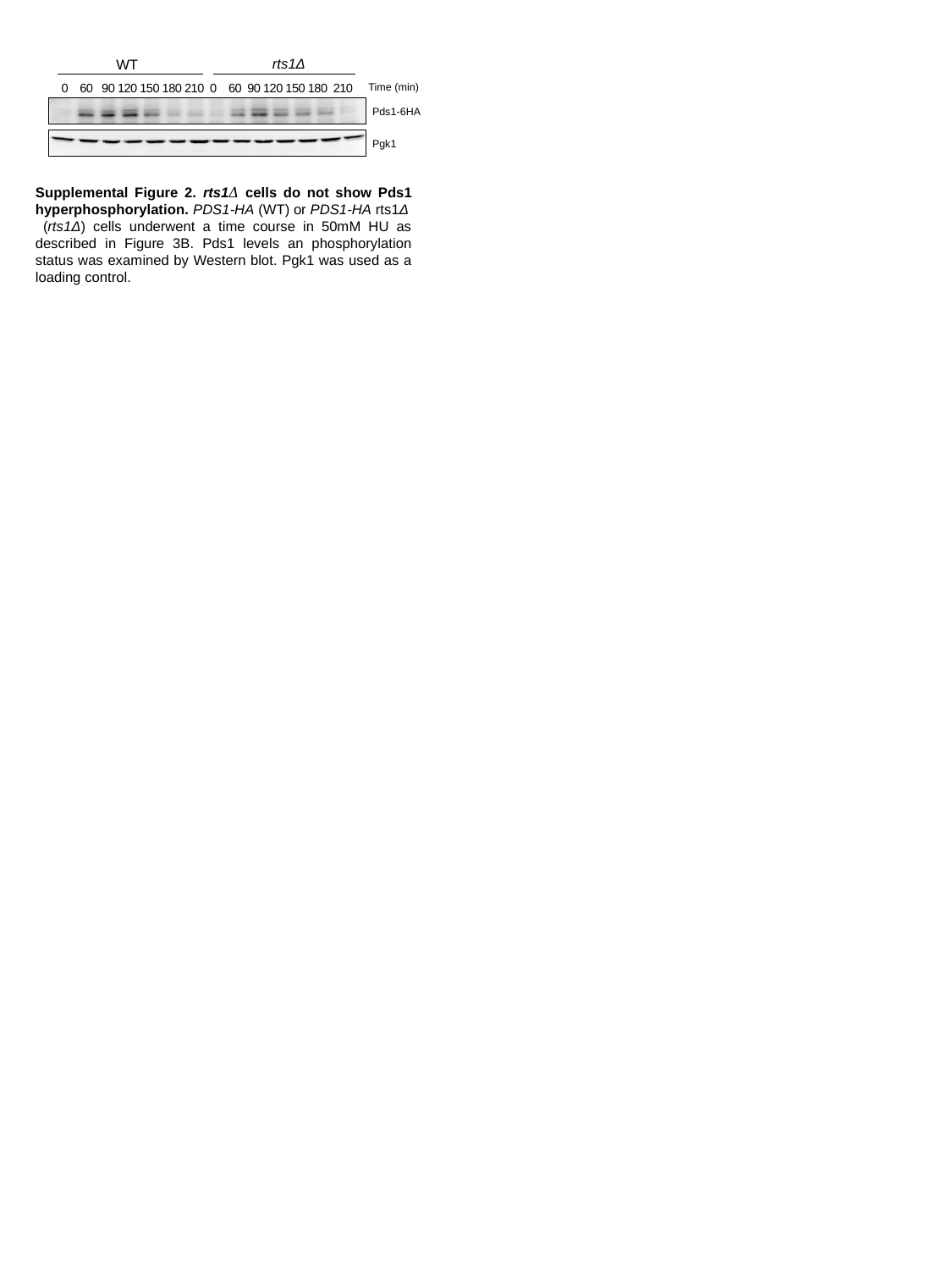

rts1Δ
WT
 0 60 90 120 150 180 210 0 60 90 120 150 180 210
Time (min)
Pds1-6HA
Pgk1
Supplemental Figure 2. rts1𝛥 cells do not show Pds1 hyperphosphorylation. PDS1-HA (WT) or PDS1-HA rts1Δ
 (rts1Δ) cells underwent a time course in 50mM HU as described in Figure 3B. Pds1 levels an phosphorylation status was examined by Western blot. Pgk1 was used as a loading control.

### Slide 3
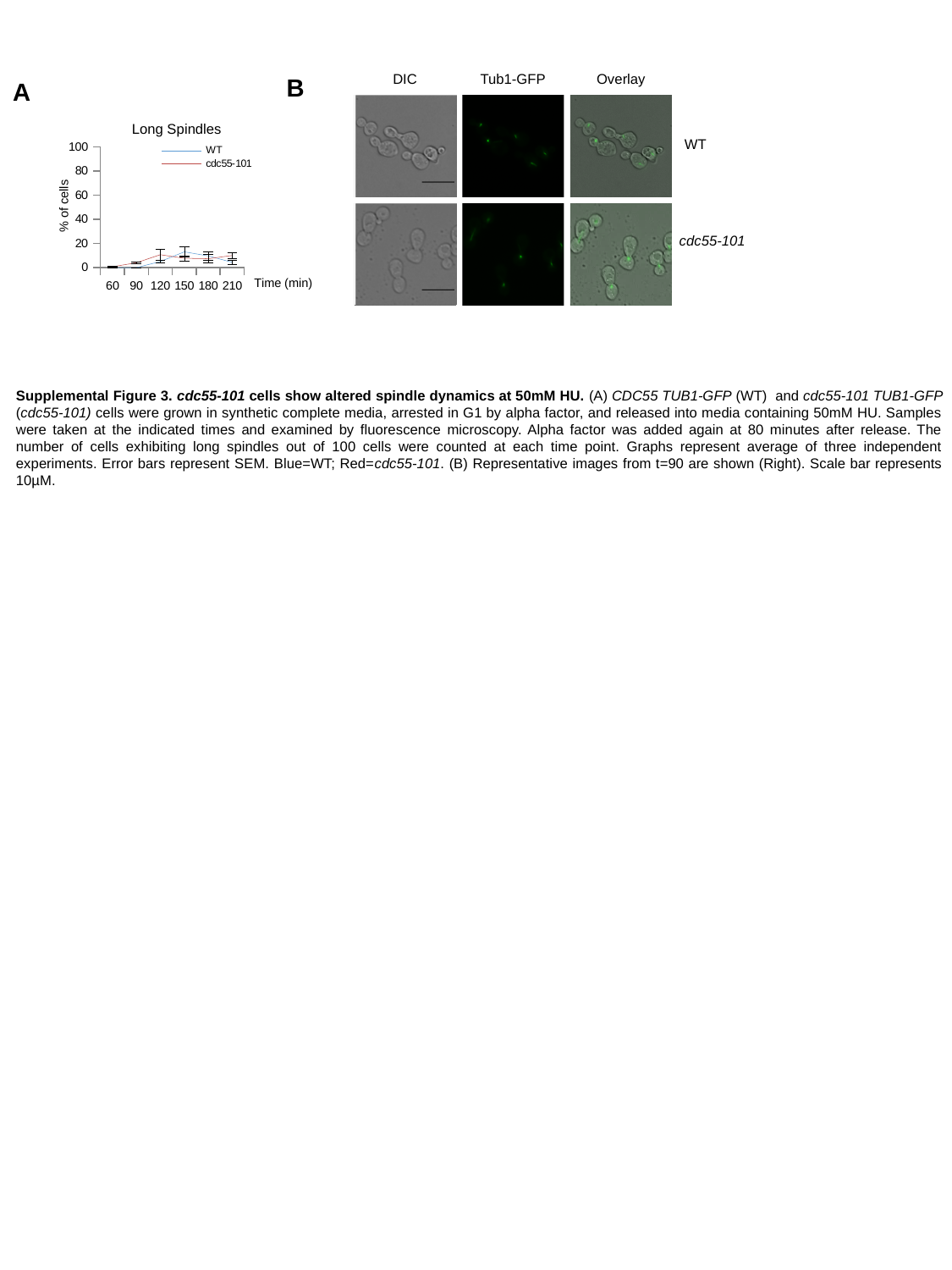

B
A
DIC
Tub1-GFP
Overlay
Long Spindles
#### Chart
| Category | WT | cdc55-101 |
|---|---|---|
| 60 | 0.0 | 0.666666666666667 |
| 90 | 0.0 | 4.0 |
| 120 | 5.0 | 10.66666666666667 |
| 150 | 13.0 | 7.666666666666667 |
| 180 | 9.66666666666667 | 7.333333333333332 |
| 210 | 4.333333333333332 | 10.0 |% of cells
Time (min)
WT
cdc55-101
Supplemental Figure 3. cdc55-101 cells show altered spindle dynamics at 50mM HU. (A) CDC55 TUB1-GFP (WT) and cdc55-101 TUB1-GFP (cdc55-101) cells were grown in synthetic complete media, arrested in G1 by alpha factor, and released into media containing 50mM HU. Samples were taken at the indicated times and examined by fluorescence microscopy. Alpha factor was added again at 80 minutes after release. The number of cells exhibiting long spindles out of 100 cells were counted at each time point. Graphs represent average of three independent experiments. Error bars represent SEM. Blue=WT; Red=cdc55-101. (B) Representative images from t=90 are shown (Right). Scale bar represents 10µM.

### Slide 4
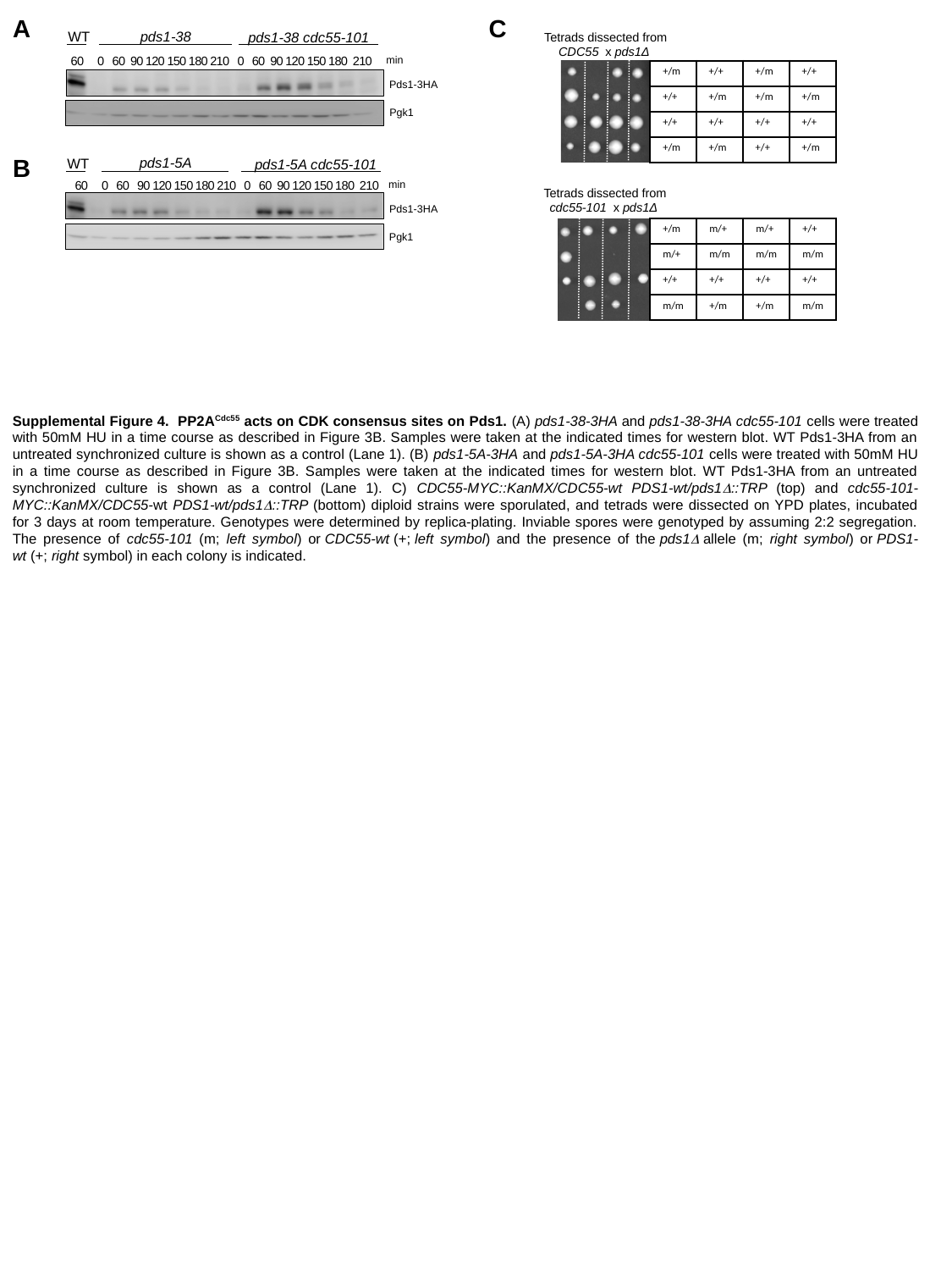

A
C
pds1-38
pds1-38 cdc55-101
 60 0 60 90 120 150 180 210 0 60 90 120 150 180 210
min
Pds1-3HA
Pgk1
WT
Tetrads dissected from CDC55 x pds1Δ
| +/m | +/+ | +/m | +/+ |
| --- | --- | --- | --- |
| +/+ | +/m | +/m | +/m |
| +/+ | +/+ | +/+ | +/+ |
| +/m | +/m | +/+ | +/m |
B
pds1-5A
WT
pds1-5A cdc55-101
min
60 0 60 90 120 150 180 210 0 60 90 120 150 180 210
Tetrads dissected from cdc55-101 x pds1Δ
Pds1-3HA
| +/m | m/+ | m/+ | +/+ |
| --- | --- | --- | --- |
| m/+ | m/m | m/m | m/m |
| +/+ | +/+ | +/+ | +/+ |
| m/m | +/m | +/m | m/m |
Pgk1
Supplemental Figure 4. PP2A^Cdc55^ acts on CDK consensus sites on Pds1. (A) pds1-38-3HA and pds1-38-3HA cdc55-101 cells were treated with 50mM HU in a time course as described in Figure 3B. Samples were taken at the indicated times for western blot. WT Pds1-3HA from an untreated synchronized culture is shown as a control (Lane 1). (B) pds1-5A-3HA and pds1-5A-3HA cdc55-101 cells were treated with 50mM HU in a time course as described in Figure 3B. Samples were taken at the indicated times for western blot. WT Pds1-3HA from an untreated synchronized culture is shown as a control (Lane 1). C) CDC55-MYC::KanMX/CDC55-wt PDS1-wt/pds1Δ::TRP (top) and cdc55-101-MYC::KanMX/CDC55-wt PDS1-wt/pds1Δ::TRP (bottom) diploid strains were sporulated, and tetrads were dissected on YPD plates, incubated for 3 days at room temperature. Genotypes were determined by replica-plating. Inviable spores were genotyped by assuming 2:2 segregation. The presence of cdc55-101 (m; left symbol) or CDC55-wt (+; left symbol) and the presence of the pds1Δ allele (m; right symbol) or PDS1-wt (+; right symbol) in each colony is indicated.
