## Supplementary material for "PP2A^Cdc55^ dephosphorylates Pds1 to inhibit spindle elongation": Table

| RUY508 | *MATa his3-11,15 leu2-3,112 trp1-1 ura3-1 can1-100* |  |
| --- | --- | --- |
| SKY029 | *MATalpha cdc55-myc::KanMX* | From S. Yoshida (VR1427) |
| SKY030 | *MATalpha cdc55-101-myc::KanMX* | From S. Yoshida (VR1821) |
| SKY177 | *Matalpha mec1:trp1::KanMX CDC55-MYC::KanMX sml1::HIS* | This Study |
| SKY178 | *Matalpha mec1:trp1::KanMX cdc55-101-MYC::KanMX sml1::HIS* | This Study |
| SKY135 | *MATa sml1::HIS3 mec1::trp1::KanMX* | From F. Cross (RUY321) |
| SKY185 | *Mata sml1::HIS* | This Study |
| SKY179 | *Mata cdc55-101-MYC::KanMX sml1::HIS* | This Study |
| SKY187 | *Mata bar1::URA CDC55-MYC::KanMX rad53::LEU sml1::HIS* | This Study |
| SKY189 | *Mata bar1::URA cdc55-101-MYC::KanMX rad53::Leu sml1::HIS* | This Study |
| SKY209 | *Mata bar1 CDC55-MYC::KanMX promURA::tetR::GFP::LEU cenIV::tetOx448::URA ADE2* | This Study |
| SKY210 | *Mata bar1 cdc55-101-MYC::KanMX promURA::tetR::GFP::LEU cenIV::tetOx448::URA ADE2* | This Study |
| SKY160 | *Mata Pds1-6xHA::TRP Cdc55-MYC::KanMX bar1* | This Study |
| SKY171 | *Mata bar1 ADE2 cdc55-101-MYC::KanMX PDS1-6xHA::TRP* | This Study |
| SKY173 | *Mata bar1 cdc55-101-MYC::KanMX Pds1-6xHA::Trp TUB1-GFP::HIS ADE2* | This Study |
| SKY205 | *Mata bar1 pds1-38-HA::URA TUB1-GFP::HIS* | This Study |
| SKY172 | *Mata bar1 CDC55-MYC::KanMX Pds1-6xHA::Trp TUB1-GFP::HIS ADE2* | This Study |
| SKY226 | *Mata bar1 CDC55-MYC::KanMX pds1-38-HA::URA* | This Study |
| SKY228 | *Mata bar1 cdc55-101::KanMX pds1-38-HA::URA* | This Study |
| SKY248 | *Mata bar1 TUB1-GFP::HIS pds1-38-3xHA::URA cdc55-101-MYC::KanMX* | This Study |
| SKY233 | *Matalpha Pds1-6xHA::TRP Esp1-9xMYC::TRP ADE2* | This Study |
| SKY234 | *Mata cdc55::KanMX Pds1-6xHA::TRP Esp1-9xMYC::TRP ADE2* | This Study |
| SKY279 | *Mata bar1 pds1::PDS1-GFP-TRP* | This Study |
| SKY280 | *Mata bar1 pds1::PDS1-GFP-TRP cdc55-101-MYC::KanMX* | This Study |
| SKY180 | *Matalpha CDC55-MYC::KanMX mad2::KanMX ADE2* | This Study |
| SKY149 | *Mata bar1 ADE2 CDC16-HA::TRP* | This Study |
| SKY197 | *Mata bar1 cdc55-101-MYC::KanMX CDC16-6xHA::TRP* | This Study |
| SKY181 | *Mata cdc55-101-MYC::KanMX mad2::KanMX ADE2* | This Study |
| SKY193 | *Mata bar1::URA cdc55-101-MYC::KanMX swe1::TRP* | This Study |
| SKY194 | *Mata bar1::URA CDC55-MYC::KanMX swe1::TRP* | This Study |
| SKY281 | *Mata bar1 ADE2 mih1𝛥::KanMX* | This Study |
| SKY285 | *Mata bar1 ADE2 CDC55-GFP-NLS::KanMX* | This Study |
| SKY287 | *Mata bar1 ADE2 CDC55-GFP-NES::KanMX* | This Study |
| SKY175 | *Mata swe1Δ::TRP1 bar1Δ::URA3* | This Study |
| SKY230 | *Mata bar1 rts1::KanMX Pds1-6xHA::TRP* | This Study |
| SKY244 | *Mata bar1 pds1-5A-3xHA* | This Study |
| SKY247 | *Mata bar1 pds1-5A-3xHA cdc55-101-MYC::KanMX* | This Study |
| SKY266 | *Mata bar1 CDC55-MYC::KanMX chk1::KanMX* | This Study |
| SKY267 | *Mata bar1 cdc55-101-MYC::KanMX chk1::KanMX* | This Study |
| SKY174 | *Mata pds1Δ::TRP1 bar1Δ::URA3* | From D. Quintana (YRP33) |
